# Phase diagrams for biophysical fitness landscape design

**DOI:** 10.1101/2025.09.10.675274

**Authors:** Vaibhav Mohanty, Eugene I. Shakhnovich

## Abstract

Viral populations often evolve toward the top of fitness landscapes by mutating their surface proteins to escape host antibodies and/or to bind more tightly to host cell receptors. As they climb fitness landscapes, newly mutated strains can spread across host populations, causing new waves of infection and death. As a pandemic preparedness strategy, we recently introduced computational protocols collectively called fitness landscape design (FLD) [Mohanty and Shakhnovich, *Proc. Natl. Acad. Sci*. (2026)], which consist of using antibody sequences and concentrations as tunable control parameters to proactively restructure the shape of a viral surface protein’s biophysical fitness landscape for optimal suppression of dangerous escape mutants. Until now, previous evidence for the feasibility of FLD has been numerical and simulated. Here, we derive an explicit analytical theory for the FLD phase diagram in the case of viral surface protein evolution in the presence of a neutralizing antibody, providing tight bounds on the designability frontiers describing the extent to which fitnesses of different protein sequences can be tuned independently of each other. We then leverage a recently published experimental dataset of binding affinities between over 62,000 antibody variants and each of three influenza surface glycoproteins to construct experimental FLD phase diagrams, which show close agreement with theory. These results support the feasibility of engineering quantitatively programmable fitness landscapes for laboratory protein evolution experiments.

## I. INTRODUCTION

Evolution of a population is often viewed as a semi-random “survival of the fittest” process in which natural selection contributes a driving force towards increased fitness over time, while genotype mutations facilitate exploration of the sequence space [1–4]. Mathematically, absolute fitness *F* (**s**) is defined as the growth rate of a strain or species **s**; i.e., *dn*(**s**)*/dt* = *F* (**s**)*n*(**s**), where *n*(**s**) is the number of individuals of type **s** [1]. First introduced by Sewall Wright more than 90 years ago [5], the mapping from genotype sequence **s** to fitness *F* (**s**) is visually represented as a mountain-like structure termed a *fitness landscape*, which has served as a useful tool for understanding evolutionary trajectories. The structure of a fitness landscape can be affected by intrinsic factors, such as protein mutations which impact organism survivability, or interactions with the environment, such as protein-protein interactions between a cell surface protein and extracellular peptides [6–13].

Viral populations also climb toward the top of fitness landscapes as they circulate within their host populations. Lytic viruses, such as influenza, norovirus, and SARS-CoV-2, primarily replicate by first attaching their surface-expressed viral attachment proteins (VAPs) [14] to a host’s cell’s surface receptor, entering the host cell, replicating within the cell, and then lysing the host cell in order to release the viral offspring. Mutations to these VAPs that lead to higher affinity to the host cell’s receptor increase the rate of viral entry into host cells, improving the VAP’s functionality, which can be considered an *intrinsic* contribution to its fitness. On the other hand, deployment of adaptive immune responses, which can be considered part of the modifiable environment of the viral population, can decrease *extrinsic* contributions to viral fitness. For example, targeted antibodies circulating in the local environment can bind to the VAPs, potentially blocking viral entry into the host cells and/or removing virions from circulation altogether, reducing the overall viral replication rate. Recent experimental and epidemiological evidence has shown that SARS-CoV-2 and norovirus fitness landscapes can be accurately predicted from statistical physics-based models of viral fitness in which a viral strain’s fitness is proportional to the Boltzmann probability of its VAP being bound to a host cell receptor while not bound to any circulating antibody [15–17]. Accurate modeling of fitness landscapes for strains differentiated by their VAP mutations can enable new and effective approaches for vaccine design and pandemic preparedness [17, 18].

Throughout this paper, we focus our discussion of evolution to specifically consider the evolution of viruses whose VAPs (which we may also call viral entry proteins, viral surface proteins, target proteins, target antigens, or surface antigens) are subject to mutation and selection pressure. In particular, we take inspiration from a particular experimental setup we have previously realized in norovirus [15] and introduce a model of *in vitro* laboratory evolution involving the competition of viral strains in the presence of host cells and a chosen neutralizing antibody; the experimental setup with the three distinct classes of proteins involved in the model are depicted in Figure 1(a). The antibodies in the broth act as an *in vitro* proxy for the adaptive immune response, preventing viral entry into host cells. Different viral strains’ fitnesses are then determined by a combination of host-VAP interaction strength (intrinsic contribution) and antibody-VAP interaction strength. With sufficient computational or experimental data for different VAP mutants, comprehensive empirical fitness landscapes for the corresponding strains can be constructed [15–17, 19].

**FIG. 1.**
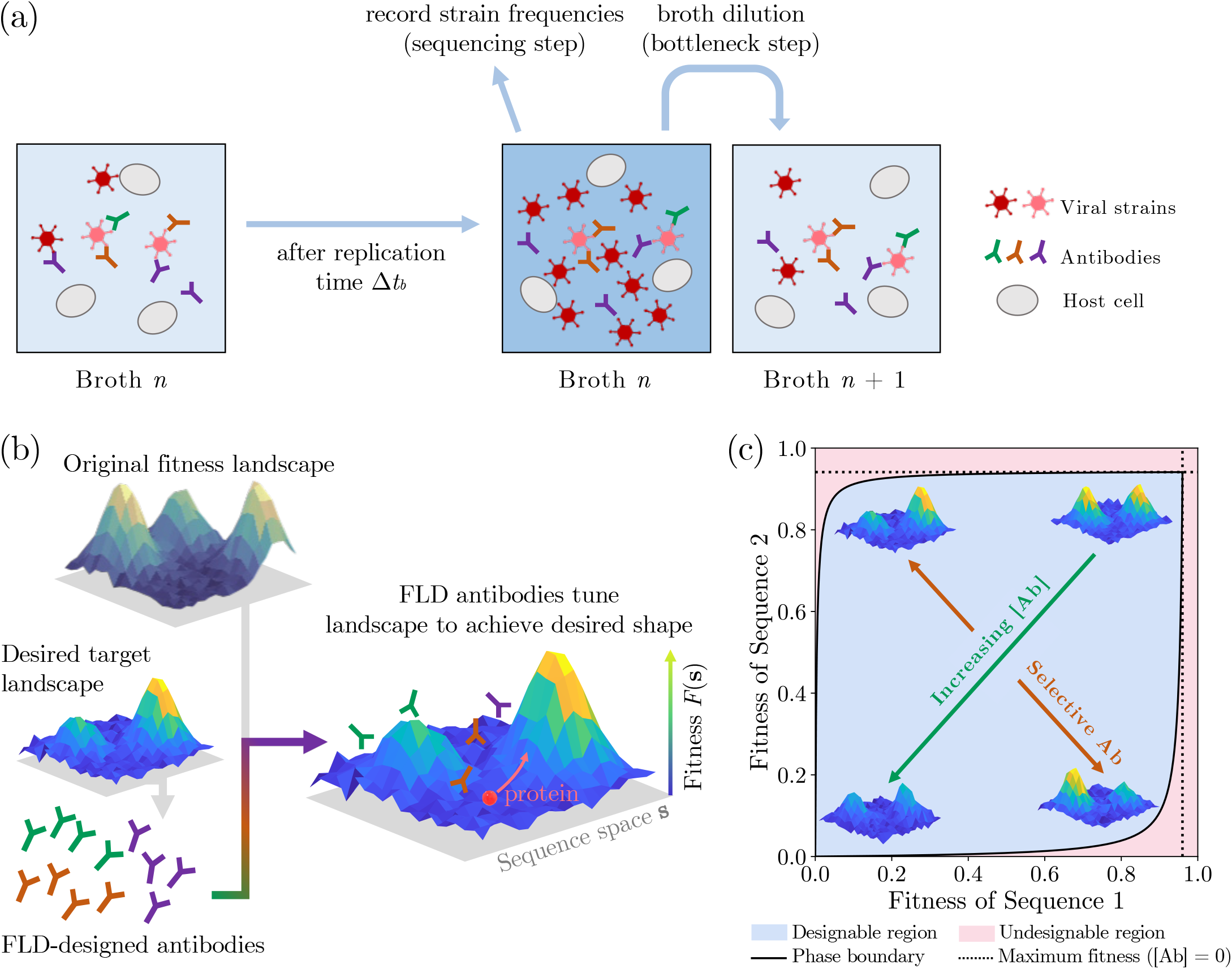
(a) Schematic depicting viral bulk culture in the presence of host cells and antibodies with serial passage. (b) Schematic for fitness landscape design, which uses antibodies used as tunable control parameters to customize the biophysical fitness landscape of a viral attachment protein by suppressing the fitnesses of different protein sequences with user-defined penalties. Also shown is a physical analogy to FLD in designing custom potential energy landscapes for ionic trapping. (c) Schematic phase diagram for FLD, showing how the fitnesses of two different viral strains with distinct viral attachment protein mutations (represented by the tops of the two peaks) can be suppressed relative to one another.

In the context of the aforementioned *in vitro* model of viral evolution (Figure 1(a)), we recently introduced *fitness landscape design* (FLD) [17], which is the task of remolding a target VAP’s biophysical fitness landscape for optimal suppression of dangerous variants by adjusting the sequences and/or concentrations of the antibodies which differentially impact the extrinsic contributions to viral strains’ fitnesses because antibodies selectively bind certain VAPs more strongly than others. Essentially, in FLD, antibodies are tunable *environmental* control parameters which selectively suppress a VAP’s biophysical fitness landscape according to a user-specified template landscape (Figure 1(b)). FLD algorithms [17] have been developed to propose or identify antibody environments which can specifically block dangerous mutants and suppress viral escape fitness trajectories, offering new strategies for biosecurity and pandemic preparedness. Some physically inspired studies have studied control of evolutionary dynamics purely from a dynamical systems perspective or by regulating drug dosage or chemical environments in real time [20–23], but FLD [17] is the first to suggest that the fitness landscape underlying evolution can itself be shaped into a quantitatively predictable structure to tackle infectious disease. As an application, we showed that our FLD algorithms can discover cross-reactive antibodies which proactively suppress the fitness trajectories of viral escape mutants [17].

FLD relies on the idea that antibody-VAP interactions have sufficient flexibility such that different viral strains can experience a wide range of possible fitnesses. For example, in the *in vitro* viral evolution experiment, in the high antibody concentration limit within the viral culture—even with weakly binding antibodies—will generally lead to low fitnesses for all strains because the vast number of antibodies will neutralize most of the viruses before they can bind to a host cell. On the other hand, having zero antibodies in solution with the viral bulk culture will lead to the maximum possible fitneses for each strain because host-VAP binding would not be competitively inhibited by antibody-VAP binding. However, numerical evidence has shown that it is possible to discover selective antibodies which can suppress some viral strains’ fitnesses but not others, allowing for many degrees of freedom in tuning fitness landscape shapes by finding appropriate antibody sequences [17]. These possibilities are described by the schematic FLD phase diagram in Figure 1(c), which characterizes what fitness landscape assignments are “designable” or “undesignable.”

In this work, we derive an explicit analytical theory for the shape of the FLD phase diagram, in agreement with previous numerical estimates [17]. We then leverage a recently published experimental dataset of over 62, 000 antibodies’ binding affinities to each of three different influenza surface antigens to construct the first experimental FLD phase diagrams, demonstrating excellent agreement with theory and supporting the notion that FLD is experimentally realizable. Finally, we discuss generalizations of FLD to multiple VAP mutants and polyclonal antibody mixtures.

## II. ANALYTICAL THEORY

### A. FLD phase diagram boundaries: pairs of viral strains with distinct VAP mutations

We now develop an analytical theory for the boundary between the *designable* and *undesignable* regions of the FLD phase diagram. For viral surface protein evolution, biophysical properties—particularly, binding affinities— of viral-host cell binding and viral-antibody are quantitatively predictive of the viral strains’ fitnesses. A statistical mechanics-based biophysical model of viral fitness which we showed can be analytically derived from chemical reaction kinetics between viral, host cell, and antibody proteins [17] aptly captures the empirical fitness data. The viral fitness can be analytically expressed as

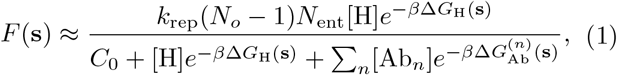

where *k*_rep_ is a virion’s microscopic rate constant for cell entry and replication, *N*_*o*_ is the average number of offspring produced in a single replication event, *N*_ent_ is the number of viral surface proteins which can potentially be used for cell entry, [H] is the total concentration of host cell receptors, [Ab_*n*_] is the total concentration of the *n*-th type of antibody, Δ*G*_H_(**s**) is the binding free energy between the viral entry protein and host cell receptor, 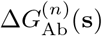 is the binding free energy between the *n*-th antibody and the entry protein of a viral strain whose VAP has sequence **s**, *β* = 1*/*(*RT* ) is proportional to the inverse temperature *T* times the ideal gas constant *R*, and *C*_0_ is a reference concentration that sets units appropriately. This model has been extensively validated by simulations of viral serial dilution experiments [17, 24], by *in vitro* growth experiments in norovirus [15], and by epidemiological infectivity data for SARS-CoV-2 [16].

To make analytical calculations tractable, we work in the case where we have two viral strains (“strain 1” and “strain 2”) with distinct VAP sequences (“sequence 1” and “sequence 2”, denoted as **s**_1_ and **s**_2_) and only one antibody whose concentration [Ab] and sequence serve as control parameters. We use rescaled fitnesses with constants *k*_rep_(*N*_*o*_ − 1)*N*_ent_ = 1 for mathematical convenience. The biophysical fitness model simplifies to

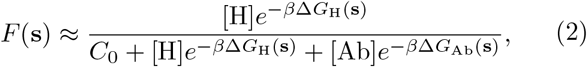

for each of the two viral strains whose VAPs are assumed to differ by mutations represented appropriately in **s**_1_ and **s**_2_. The fitness of viral strain 1 will eb represented by the shorthand *F*_1_ ≡ *F* (**s**_1_), and the fitness of strain 2 will be represented by the shorthand *F*_2_ ≡ *F* (**s**_2_).

To find the boundaries of the designable region, we ask what are the maximum and minimum strain 2 fitnesses that can be realized for a given value of strain 1 fitness? To calculate the bounds on *F*_2_ given *F*_1_, we first rearrange eq. (2)

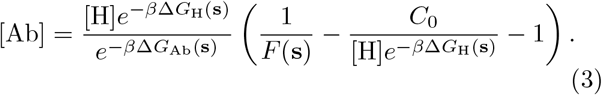

The above equation holds separately for strains 1 and 2, which would both approximately experience the same antibody concentration in the limit where the antibody concentration great exceeds viral strain concentrations, which is a clinically relevant limit in viral infections such as SARS-CoV-2 in vaccinated individuals [17]. Equating the antibody concentrations gives us

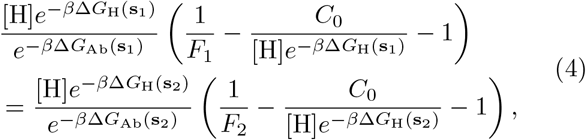

and solving for *F*_2_ in terms of *F*_1_, we have

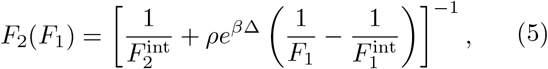

where we have defined 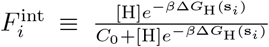 as the “intrinsic” fitness of the viral strain in the *in vitro* experiment (Figure 1(a)) with *zero* antibody concentration, Δ ≡ Δ*G*_Ab_(**s**_1_) − Δ*G*_Ab_(**s**_2_) as the free energy difference between the two viral strains bound states to the antibody, and 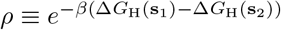, which is equivalent to the ratio of the two VAP mutants’ dissociation constants with the host protein.

Now, we note that host-viral binding affinities Δ*G*_H_(**s**_1_) and Δ*G*_H_(**s**_2_) are fixed (because we cannot manually mutate the host receptor protein), but we do have control over the choice of antibody, which is chosen from a library of *M* antibodies. We work within the approximation that the two antibody-antigen binding free energies Δ*G*_Ab_(**s**_1_) and Δ*G*_Ab_(**s**_2_) are chosen from a bivariate Gaussian distribution:

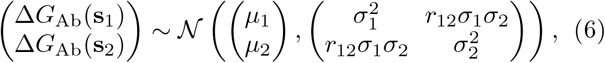

where means *µ*_1_ and *µ*_2_, standard deviations *σ*_1_ and *σ*_2_, and the Pearson correlation *r*_12_ ∈ [ ™1, 1] are taken to be be parameters fixed by the choice of the two target antigens **s**_1_ and **s**_2_. The antibody we choose is taken from a finite library of size *M* ; for instance, if we consider all combinatorial amino acid variation at *L* paratope sites, *M* = 20^*L*^. Like in Derrida’s random energy model (REM) [25, 26], having a finite (even if a large) number of possible samples means that there will be a finite ground state energy. The frontiers of the designable phase will be determined by such extremal antibody samples from the library which maximize the magnitude Δ = Δ*G*_Ab_(**s**_1_) Δ*G*_Ab_(**s**_2_), the difference between the two antibody-antigen binding free energies. Such antibodies are selective in that they tightly bind to one VAP mutant but not the other, causing one viral strain to have low fitness while the other is high.

From the Gaussian approximation, we know that Δ is also Gaussian distributed: Δ ∼ *N* (*µ*_Δ_, *σ*_Δ_), where we define *µ*_Δ_ ≡ *µ*_1_ − *µ*_2_ and 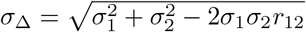. A standard result in classical extreme-value statistics [27] provides an approximation to the mean of the maximum (or minimum) value of Δ in the limit of large *M* samples:

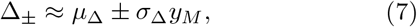

where

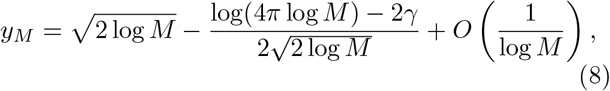

and *γ* is the Euler-Mascheroni constant. The leading order contribution to *y*_*M*_ is exactly the ground state energy of the REM [26], and the sub-leading order term which scales as 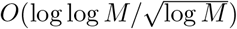 is related to finite-size corrections to the ground state energy of the REM [28]. Heuristically, we can understand that the Gaussian approximation employed here works because the antibody-antigen complex in the bound state can be treated as a heteropolymer in a folded state, with the variable monomers (or spins, in the REM analogy) consisting of *L* amino acids on the antibody surface. The equivalence of the spin-glass model of folded heteropolymers to the REM is well-known result in protein folding physics [29]. The antibody for which Δ = Δ_−_ is the “ground state” antibody which selectively binds VAP mutant 1 more strongly than it binds to VAP mutant 2, compared to any other antibody in the library. Conversely, the Δ = Δ_+_ solution yields an antibody which selectively binds VAP mutant 2 more strongly than it binds to VAP mutant 1, compared to any other antibody in the library.

We can now calculate maximum possible strain 2 fitness 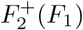 given a value of strain 1 fitness and minimum possible strain 2 fitness 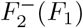 given a strain 1 fitness; these will be obtained from the Δ = Δ_−_ and Δ = Δ_+_ antibodies, respectively

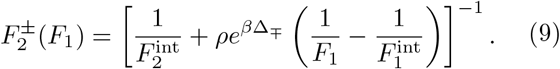

These designability boundaries—a central result of this work—are valid on the strain 1 fitness domain 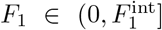, which corresponds to the strain 2 fitness range 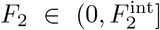. The exact same boundaries could be obtained by repeating the derivation but instead maximizing or minimizing *F*_1_ for fixed *F*_2_, yielding relabeled but equivalent curves 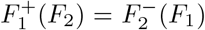 and 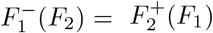.

We plot theoretical phase diagrams using eq. (9) in Figure 2 across a range of antibody library sizes *M* and Pearson correlations *r*_12_. The designable region (blue) is generally an asymmetric leaf-shaped region within the unit square whose sharp corners are placed at (0, 0) and 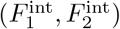. The two curves 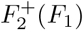 and 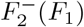 (black solid lines) respectively outline the top and bottom borders of the designable region, and they converge at two points, representing limiting cases of the antibody concentration: in the limit of no antibodies ([Ab] = 0), both fitnesses adopt their intrinsic limits 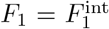 and 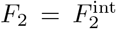 (black dotted lines). In the context of this mathematical fitness model, fitnesses above these intrinsic limits are unobtainable since they are reached in the limit [Ab] → 0, and antibody concentration cannot go below this limit. In the limit of high antibody concentration ([Ab] → ∞), we see from the original fitness equation eq. (2) that both fitnesses *F*_1_, *F*_2_ → 0. Outside of the designable region is the undesignable region (red), where fitness assignments are unreachable by any choice of antibody. Although we work in the single antibody limit, we can also obtain the estimated fitness curve (purple dashed line) for a large random ensemble of different antibodies with uniform concentrations by taking the annealed approximation 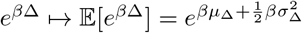 within eq. (5).

**FIG. 2.**
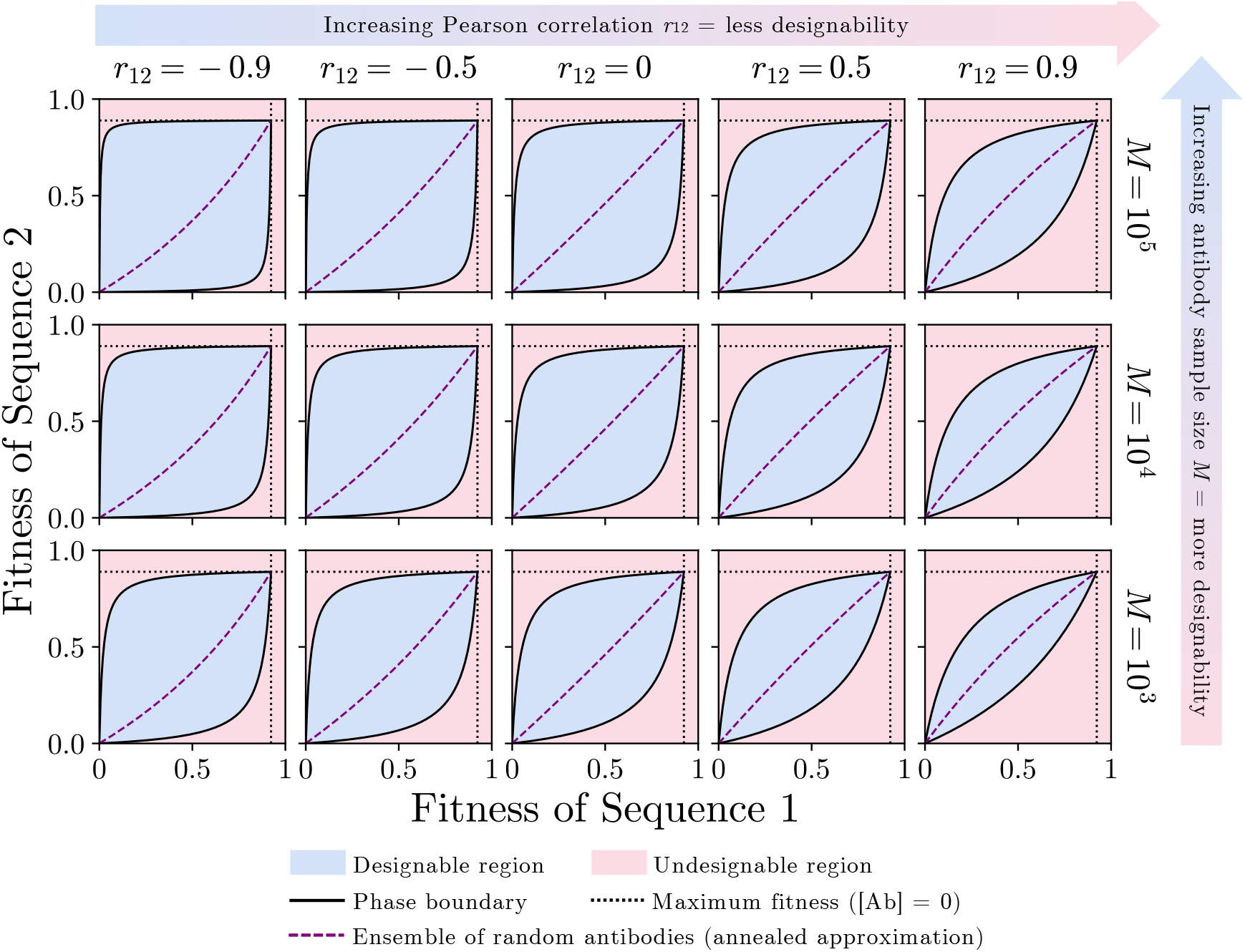
Example theoretical FLD phase diagrams from eq. (9). The area of the designable region expands with decreasing Pearson correlation *r*12 and with increasing antibody library size *M* .

We see in Figure 2 that the designable region expands with increasing antibody library size *M* and with decreasing Pearson correlation *r*_12_ between Δ*G*_Ab_(**s**_1_) and Δ*G*_Ab_(**s**_2_). Increasing *M* also increases the expected maximum/minimum of the free energy samples from the Gaussian distribution in eq. (6). The latter trend makes sense when we contextualize the biophysical meaning of *r*_12_. Two VAP mutants have high *r*_12_ when they bind with similar free energy to the same antibody variant. If the binding free energies are likely to be similar, so both strains’ fitnesses are likely to be similar, which decreases the area of the designable region. The Pearson correlation *r*_12_ may not necessarily be interpreted simply as a measure of chemical similarity between the two strains’ VAP sequences because many sequences can give the same fold [30]. Instead, we should think of *r*_12_ as how well a population of antibodies discriminates between two antigen sequences **s**_1_ and **s**_2_. We now develop an analytical theory to quantitatively explain this behavior of the designable region.

### B. Codesignability score: FLD phase diagram area

The area of the designable region relative to the (trivial) area of the unit square is called the *codesignability score* CDS(**s**_1_, **s**_2_) for the two strains, as defined and computed numerically in ref. [17]. High codesignability score indicates increased flexibility for fitness assignment: an antibody can be found such that strain 1 fitness can be high while strain 2 fitness is low—or vice versa—and it is possible to modulate antibody concentration so that the fitnesses are both high or both low. We previously showed that codesignability scores for all pairs of strains in a larger set are useful for understanding what groups of strain sequences may have their fitnesses designed together, though all of our computations of codesignability scores were numerical [17]. Here, we can compute codesignability score analytically.

By definition, codesignability score is the area of the designable region, which is computed by finding the area between the 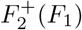 and 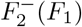 curves:

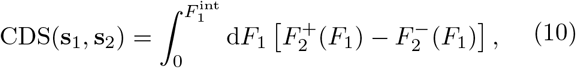

which has an exact integral solution:

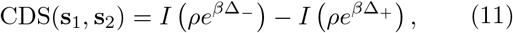

where

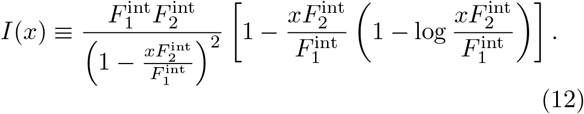

We can also prove that codesignability score monotonically decreases with Pearson correlation *r*_12_. First, we note

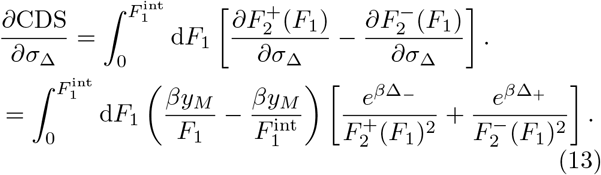

Since 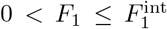, the term within the parentheses is positive. Since exponentials in the numerators and quadratic terms in the denominators are also positive, the entire integrand is positive, which means *∂*CDS*/∂σ*_Δ_ *>* 0. Noting that *∂σ*_Δ_*/∂r*_12_ = ™*σ*_1_*σ*_2_*/σ*_Δ_ *<* 0, it follows from chain rule that

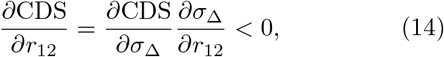

which completes the proof. The result indicates that if binding free energy distributions are positively correlated for a pair of antigens, the codesignability score decreases; in fact, the relationship between CDS and the binding free energy distribution correlation is monotonically decreasing.

## III. EXPERIMENTAL DESIGNABILITY PHASE DIAGRAMS FOR INFLUENZA ANTIGENS

### A. Experimental FLD phase diagrams, and comparison to analytical theory

The phase diagrams from theory qualitatively match the results from our original numerical FLD paper [17], where we used force field-based binding energies and support vector machines to construct simulated phase diagrams. We now show that the analytical theory also agrees with *in vitro* experimental results.

Philips et al. [31] report experimental *in vitro* log dissociation constants between *M* = 2^16^ = 65, 536 mutational variants of the human CH65 antibody and each of three mutational variants of the influenza H1 glycoprotein antigen. They chose *L* = 16 amino acid sites which varied between the unmutated common ancestral antibody and the affinity-matured CH65 antibody and performed all combinatorial mutations between those two antibody types, meaning two possible amino acids at each of the 16 sites were tested. Their binding affinity measurements, obtained using the Tite-Seq technique [32], are reported as log-dissociation constants 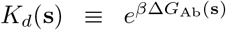, where **s** is the antigen sequence label referring to one of three influenza strains: A/Massachusetts/1/1990 (MA90), A/Solomon Islands/3/2006 (SI06), and A/Massachusetts/1/1990 with the G189E mutation (G189E). Non-binding sequences were assigned the titration boundary value ™log *K*_*d*_ = 6. After filtering out poor quality measurements, their published dataset contained collectively over 193, 000 binding affinities across the three antigens and all antibodies.

Here, we use ref. [31]’s rich dataset of reported binding affinities to construct the first-ever experimental FLD phase diagrams for each pair of influenza antigens, allowing direct comparison to our analytical calculations. Since host-antigen binding affinity measurements are unavailable, we rescale the rectangular region bounded by the dotted lines in Figure 2 to the unit square by making the replacements 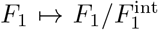 and 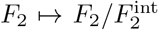, which renormalizes the fitnesses of the two antigens to the range (0, 1]. For each tested antibody, and for each pair of influenza antigens, we use the experimental binding dissociation constants from [31] to calculate Δ and substitute it into eq. (5), which is plotted on the unit square. Theoretical phase diagram boundaries are calculated by computing the means, standard deviations, and Pearson correlations of the two-antigen (Δ*G*_Ab_(**s**_1_), Δ*G*_Ab_(**s**_2_)) joint distributions and substituting into eq. (9). These values and distributions are reported Figures S1, and S2.

Pearson correlations are all positive and between 0.6 and 0.95.

These experimental results and theoretical boundaries are plotted in Figure 3. The solid lines of various colors are the transformed experimental data, while the black dashed lines indicate the theoretical boundaries separating the predicted designable (blue) and undesignable (red) regions. Figure 3(a) shows that the fitness of SI06 is generally higher than the fitness of MA90, with the designable region spanning most of the upper triangle above the central diagonal of the unit square. Figure 3(b) and (c) have their designable regions primarily spanning the lower triangle of the unit square, with SI06 generally having equal or higher fitness than G189E, which tends to have equal or higher fitness than MA90. Notably, each of the three pairs of antigens showcase one or more prominent “outlier” antibodies: orange top line in Figure 3(a), green and brown top lines as well as gray bottom line in Figure 3(b), and the pink top line in Figure 3(b). These antibodies push the designability frontier by being more selective for the antigen which generally binds more weakly to the other antibody variants. A notable feature of all three figure panels is the tightness of the theoretical bounds from eq. (9) near these outlier antibodies.

**FIG. 3.**
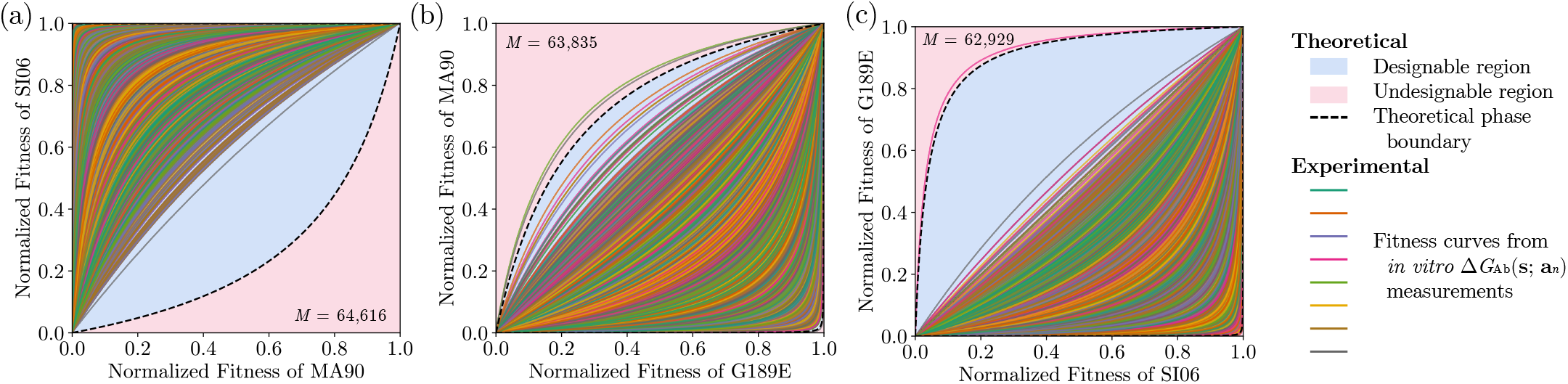
Experimental FLD phase diagrams from experimental log dissociation constants for over 62,000 antibodies with three influenza glycoprotein antigens, and theoretical phase boundaries (black, dashed) from the Gaussian approximation eq. (9). Empirical measurements are shown as solid, colorful lines.

The lower bound in Figure 3(a) and the upper bound in Figure 3(b) appear overly optimistic. This is likely due to the non-Gaussian distribution of the log dissociation constants in ref. [31]’s dataset. One contributing factor to non-Gaussianity is the assignment of ™log *K*_*d*_ = 6 to all antibody-antigen pairs which bind too weakly to be measured using Tite-Seq [31]. Another likely factor is the effect of evolutionary selection on the available antigens and antibodies. MA90 circulated in 1990 while SI06 circulated in 2006; its fitness may be expected to be higher than MA90 for most antibodies in the repertoire because it may indeed have evolved to escape related antibodies [31]. Similarly, G189E was specifically evolved in the laboratory to escape the universal common ancestral antibody, so its affinity to most of the antibodies in the repertoire are expected to be lower [31]. The extreme value statistics/REM results used in our theory rely on the Δ*G*_Ab_(**s**) samples being unbiased, i.i.d. draws from the Gaussian distribution. Since both evolutionary selection and experimental choice of amino acids played a role in the construction of the antibody library, ref. [31]’s dataset is not expected to be Gaussian.

### B. Scalability of FLD phase diagram estimation with generalized Pareto distribution tail-fitting

To address non-Gaussianity, we also construct semi-empirical phase diagrams by fitting the tails of the distribution for the free energy difference, Δ = Δ*G*_Ab_(**s**_1_) − Δ*G*_Ab_(**s**_2_), for each pair of antibodies. In extreme value theory, only the tails of the distribution and the sample size (i.e. antibody library size *M* ) affect the expected extrema. The theoretical phase diagrams from the Gaussian distribution are constructed by fitting only two parameters: the mean *µ*_Δ_ and standard deviation *σ*_Δ_. In Supplementary Appendix A, we show that by fitting each upper and lower tail using the generalized Pareto distribution (GPD) with three parameters per tail—still small compared to the number of samples—we can capture the empirically observed phase boundaries nearly perfectly (Supplementary Appendix A; Figure S5). This is expected, since the GPD fits are influenced by extremal values in the data distribution.

However, we find remarkably that even after massively downsampling the antibody dataset by ∼90%, fitting the tails to the subsampled distribution is still predictive of the phase boundaries for the full antibody dataset containing over 62,000 antibodies, as shown in Figure 4 (see Supplementary Appendix A for step-by-step construction). We downsampled the complete antibody dataset to 10% of its original size in 10,000 independent trials and computed the phase boundaries. Figure 4 shows 25th, 50th (median), and 75th percentiles for these phase boundaries aggregated over the 10,000 trials. All predicted median phase boundaries (Figure 4: black, dashed lines) are tighter than the ones from the Gaussian approximation in Figure 3, despite having been constructed from roughly 10% of the original data. Similar analysis, but with downsampling of the original data by an additional order of magnitude to only 1%, is shown in the phase diagrams in Figure S9, which shows comparable medians to the 10% phase diagrams but broader interquartile ranges.

**FIG. 4.**
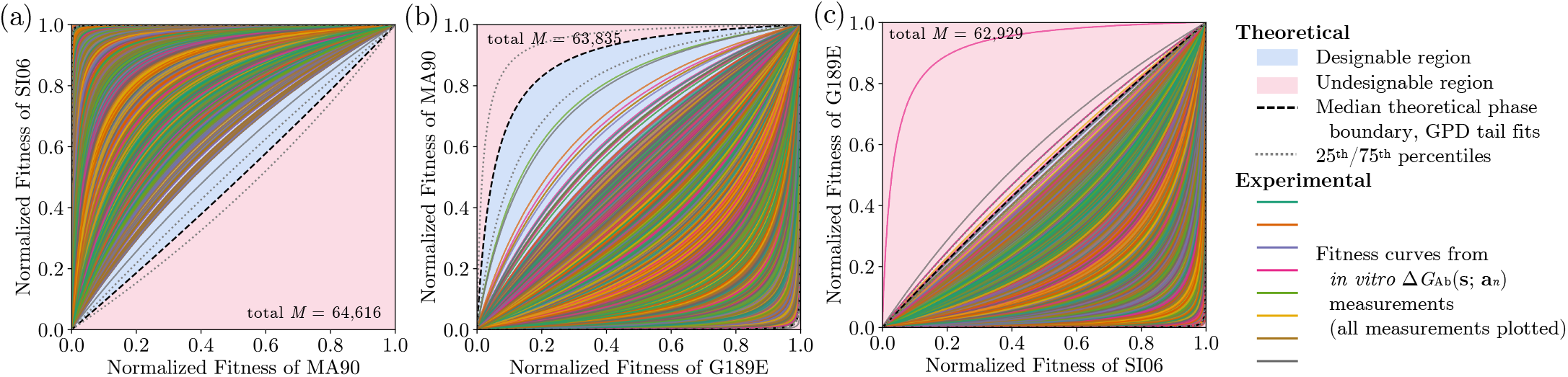
Experimental FLD phase diagrams from experimental log dissociation constants of over 62,000 antibodies with three influenza glycoprotein antigens, and semi-empiricial phase boundaries from fitting the tails of downsampled antibody datasets which are approximately 10% of the size of the original dataset. Empirical measurements are shown as solid, colorful lines. Estimated median phase boundaries from 10,000 independent downsampling trials are shown as black, dashed lines, and 25th/75th percentile phase boundaries are shown as gray, dotted lines.

Although Figure 4 and Figure S9 robustly capture phase diagram boundaries in general, any extreme value prediction which relies on a smaller set of finite samples is less likely to capture outliers which may strongly impact the GPD fits, as seen for example in Figure 4(c), where the pink antibody outlier is not captured by the phase boundaries from the downsampled fits. The quartile phase boundaries (Figure 4: gray, dotted lines) show that some boundary predictions are better than others, depending on the presence/absence of outliers drawn from the antibody library (see Figure S8 for complete distributions of expected extrema from the 10,000 trials). Using larger antibody libraries for tail fitting fixes these discrepancies, as mentioned previously (Supplementary Appendix A; Figure S5). Overall, the downsampling approach shows promise for laboratory scalability of FLD phase diagram construction by reducing the number of antibodies needed to calculate boundaries, provided that tails are captured aptly.

Finally, we emphasize that the phase diagrams computed here are limited in that completely random antibody sequence sampling was not conducted, and only around 2^16^ ∼ 6.6 × 10^4^ out of 20^16^ ∼ 6.6 × 10^20^ possible antibody mutations at the *L* = 16 sites were explored, so broader amino acid variation as well as expansion of the paratope sites may yield expanded designable (blue) regions. Also, we add that eq. (1) suggests that apparent “gaps” between the outlier antibodies and the bulk of the distribution (e.g. between the pink outlier and bulk in Figure 4(c)) may be explored by using combinations of antibodies at different concentrations—an idea supported by our random antibody ensemble sampling results in the original FLD paper [17].

Overall, we can conclude that the tail-fitting approach introduced here enables *scalability* of phase boundary prediction from a smaller set of experimental log dissociation constant measurements.

## IV. GENERALIZATION TO ADDITIONAL ANTIGENS AND POLYCLONAL MIXTURES OF ANTIBODIES

Thus far in this work, we have restricted our analysis to two viral strains’ VAPs (antigens) evolving in the presence of one antibody which has been specially chosen from a random ensemble with known properties. Accordingly, the theoretical phase diagrams we have shown have all represented, for a fixed ensemble size *M*, the fitness assignments to the two antigens that may be possible under the assumption that the appropriate *monoclonal* antibody can be extracted from the random ensemble.

Experimental phase diagrams using influenza antibodies have shown that antibody sequence variation indeed allows for a wide designable region.

But, what happens in the case where we have a *polyclonal* mixture of antibodies *simultaneously* present in the viral culture? And what if there are more than two viral strains? To address these possibilities, we depart from the random ensemble of antibodies with varied sequences and instead assume that we have a polyclonal mixture of *fixed* antibodies simultaneously present in our viral culture whose concentrations, on the other hand, are fully tunable. Thus, we depart from using extreme value statistics to estimate the shape and size of the designable region of the FLD phase diagram and instead study the geometry of the region of assignable fitnesses in the presence of a small number of antibodies whose sequences, and therefore binding free energies, are fixed. First, suppose that we study the previously considered system, with two antigens and only one antibody (i.e. not yet a polyclonal mixture of antibodies). From eq. (2), we can write the fitnesses of the two viral strains, which are defined by VAP sequences **s**_1_ and **s**_2_:

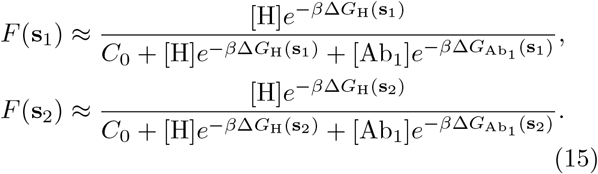

This time, the binding free energies are all fixed, so the only tunable parameter is the antibody concentration [Ab_1_]. By sweeping across the antibody concentration [Ab_1_] ∈ [0, ∞), the two above equations define a one-dimensional parametric curve in the two-dimensional fitness assignment space for viral strains 1 and 2, as shown in Figure 5(a). Such a curve is equivalent to one of the experimental curves shown in Figures 3, 4, S5, and S9. One antibody with fixed sequence can only generate a one-dimensional curve in the two-dimensional fitness assignment space.

**FIG. 5.**
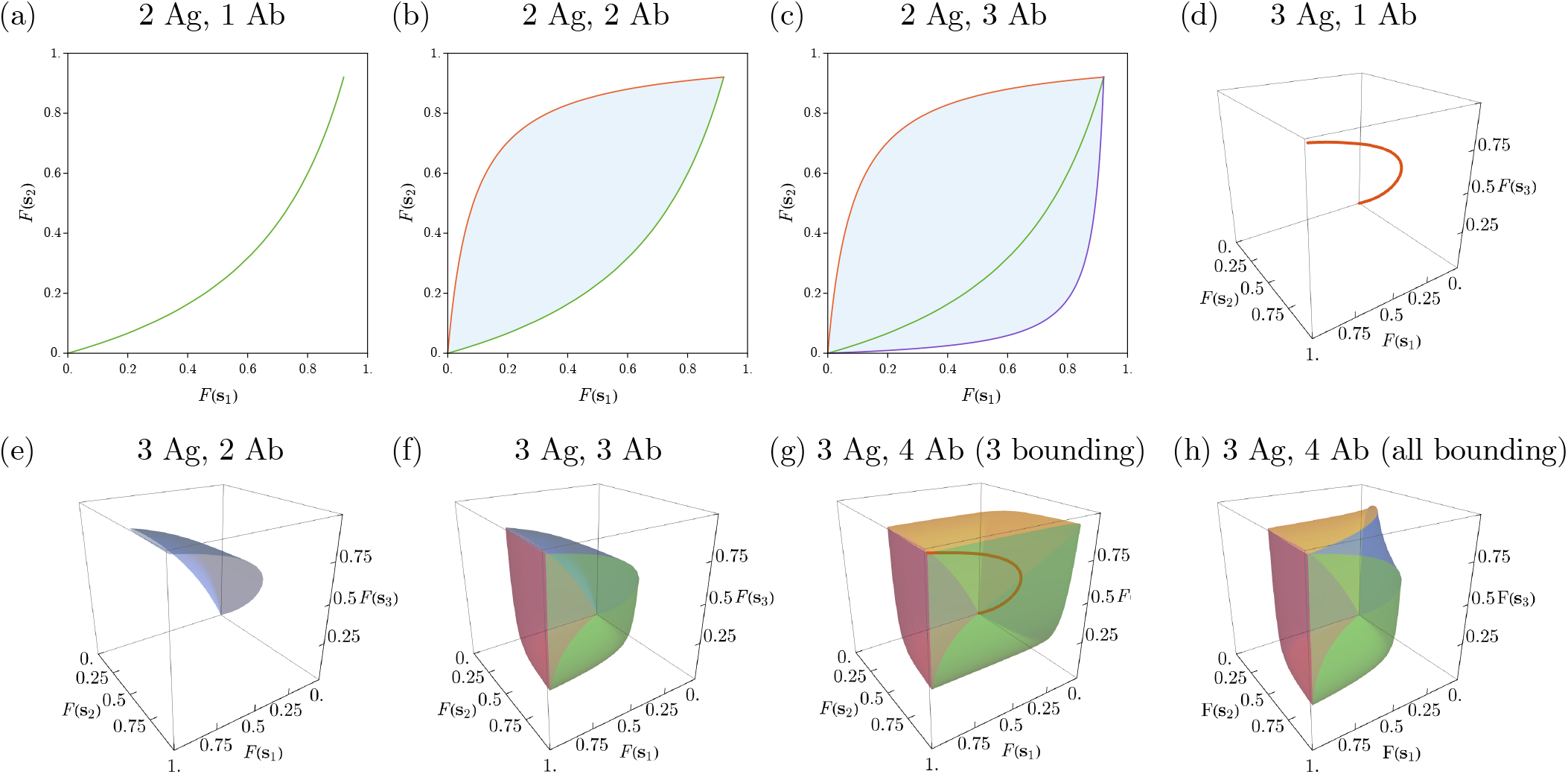
Theoretically accessible fitness assignments for different numbers of antigens and polyclonal mixture of antibodies with fixed sequences and tunable concentrations. Two-dimensional phase diagrams represent two antigens and three-dimensional phase diagrams represent three antigens. (a) One antibody generates a one-dimensional curve in the two-dimensional fitness assignment space. (b) With two antibodies (red and green) the enclosed region (blue) becomes accessible by setting the concentrations of the two antibodies appropriately. (c) A third antibody (purple) can expand designable region, but the antibody whose curve is in the interior (green) now is no longer needed to achieve fitness assignments within the designable (blue) region, as any fitness assignments within the enclosed region can be achieved with only a combination of red and purple antibodies. (d) When considering three antigens, one antibody generates a curve in the three-dimensional fitness assignment space. (e) A mixture of two antibodies creates a surface of accessible fitness assignments for the three antigens. (f) A mixture of three antibodies creates a volume of accessible fitness assignments of the three antibodies; any point within the enclosed volume can be obtained by adjusting the concentrations of the three antibodies. (g) Adding a fourth antibody can expand the volume of accessible fitness assignments; in this case, the antibody whose curve is within the interior (orange) is not needed because any fitness assignment within the now-larger region can be obtained with only three antibodies. (h) Four antibodies can, however, be chosen such that all participate in bounding the accessible region.

With two antibodies simultaneously present in the environment, the general biophysical fitness formula eq. (1) becomes

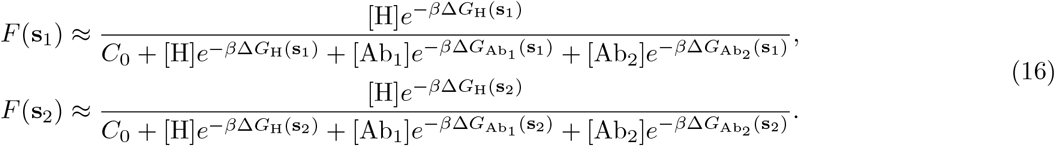

The key result is that the two fixed antibodies’ concentrations ([Ab_1_], [Ab_2_]) ∈ [0, ∞)^2^ can be swept to define a *two-dimensional region* in the fitness assignment space. In Figure 5(b), the red and green cures individually represent fitness assignment possibilities if only one or the other monoclonal antibody were present in the mixture. But, with a polyclonal mixture of the two antibodies, any fitness assignment in the region (blue in the figure) bounded by the two curves is accessible by finding the appropriate concentrations of the two antibodies. Adding a third antibody (purple in Figure 5(c)) to the polyclonal mixture can expand this region, but it also means that whichever of the three antibodies’ curves is within the interior of the accessible region is redundant, since only the antibodies corresponding to the outer two curves (red and purple in Figure 5(c)) are sufficient to produce any fitness assignment within the region.

This analysis extends to FLD for more than two antigens as well. For three antigens, we can visualize the (scaled) fitnesses in the unit cube, generalizing the unit square thus far used throughout this paper. Now, a monoclonal antibody with fixed sequence and tunable concentration generates a one-dimensional curve in the three-dimensional fitness assignment space (Figure 5(d)). A polyclonal mixture of two antibodies with fixed sequences creates a two-dimensional *surface* of accessible fitness assignments, as shown in Figure 5(e). Adding a third antibody to the polyclonal mixture creates a three-dimensional volume of accessible fitness assignments where each of the bounding surfaces are generated by the three possible pairs of antibodies (Figure 5(f)). A fourth antibody has the potential to expand the volume of accessible fitness assignments, as shown in Figure 5(g). If one of the one-dimensional antibody curves (e.g. orange in Figure 5(g)) appears within the interior of the three-dimensional volume formed by the other three antibodies, it can be excluded from the polyclonal mixture because the other three antibodies are sufficient to obtain any fitness assignment within the three-dimensional volume. However, is also possible for the four antibodies to all contribute to the bounding volume without redundancy, as shown in Figure 5(h).

We expect these principles can be generalized to a larger number *N* of FLD antigens (i.e. VAPs with unique sequences) and more complex polyclonal mixtures of antibodies. In general, we expect that it should be possible to maximize the accessible region (which is an *N* - dimensional hypervolume) for *N* VAP mutants if at least *N* antibodies are combined in a polyclonal mixture and if each of those *N* antibodies is selective for one VAP mutant each (and not cross reactive to other VAP mutants). Thus, we argue that while disease treatment can benefit from having fewer, cross-reactive broadly neutralizing antibodies, in an *in vitro* evolution setting the space of achievable fitness landscapes is expanded by using a polyclonal mixture of *narrowly* neutralizing antibodies.

## V. DISCUSSION

In summary, we have introduced analytical foundations for biophysical FLD to customize the evolutionary fitnesses of different strains with distinct VAP sequences, grounding our numerical work [17] in a theoretical basis. By working in the two-strain, one-antibody case, we have found analytically tractable closed-form expressions for the boundaries of the designable region of the designability phase diagrams. We quantitatively justified the intuition that the area of the designable region—the codesignability score—should increase when the antibody-antigen binding free energy distributions for different antigens are decorrelated. We validated the theoretical bounds on the designability phase diagrams by employing experimental *in vitro* binding affinities for over 62, 000 antibodies across three influenza antigens. Our theoretical bounds demonstrated successful ability to capture outlier antibodies which are responsible for expanding the designability frontier. We also showed that the theoretical bounds could be improved upon by fitting the tails of the free energy difference distributions directly, even when the antibody dataset was downsampled by 90%. Finally, we discussed extensions to additional antigens and polyclonal antibody mixtures.

FLD is now building a solid experimental foundation, though there are current limitations and many directions for future efforts. In recent years, biophysical fitness formulas of the form in eq. (1) used as the basis of FLD have been validated *in vitro* [15] and with epidemiological data [16]. Now, the present work uses experimental log dissociation constant measurements [31] to support the notion that the space of all possible fitness landscapes separates into designable and undesignable regions, enabling the design of custom fitness landscapes for protein evolution—an idea we first proposed in ref. [17]. Although experimental binding affinities are substituted into an experimentally validated fitness formula, a limitation of this study is that true end-to-end experimental validation of FLD should involve performing direct fitness measurements of different strains in bulk culture in the presence of FLD-designed antibodies. This is a focus of our ongoing work.

Biophysical FLD opens the door not only to improved pandemic preparedness and biosecurity via proactive vaccine design [17] by discovering proactive antibodies that simultaneously inhibit wildtype and escape antigens, but also to possible cancer therapeutics such as engineered CAR-T cells or small peptides which may prevent immune escape of cancer cells. In general, beyond viral evolution, FLD has the potential to generalize to cellular proteins, both on the surface and in the interior. In the way that antibodies dictate the extrinsic impact on fitness of a viral strain via antibody-VAP ineractions, many cell signaling pathways essential to proliferation rely on the strength of protein-protein interactions. By mutating some “protein *A*,” we may be able to control its binding strength to “protein *B*” and ultimately impact fitness should we be able to quantitatively model how *A*–*B* interactions impact proliferation rate. Since experimental evolution of cellular organisms can be carried out by selecting for fitness proxies such as surface protein fluorescence using *in vitro* compartmentalization and/or cell sorting [33–36], the designable fitness need not be proliferation rate. Lastly, widespread use of phages for directed evolution [37–39] is also an active target for applications of FLD; we suggest that FLD principles may be used for building experimental toolkits to engineer custom fitness landscape environments for experimental evolution and fundamental population genetics research.

## VI. DATA AVAILABILITY

The experimental influenza antibody-antigen data that support the findings of this article are openly available from Phillips et al., eLife, 2023 [31].

## VII. ACKNOWLEDGMENTS

This work was supported by a Hertz Foundation Fellowship (to V.M.) and by award T32GM14427 from the National Institute of General Medical Sciences (to Harvard/MIT MD-PhD Program). This work used the Accelerating Computing for Emerging Sciences (ACES) computing cluster at Texas A&M University through allocation number BIO250134 (to V.M.) from the NSF Advanced Cyberinfrastructure Coordination Ecosystem: Services & Support (NSF ACCESS) program, which is supported by U.S. National Science Foundation grants #2138259, #2138286, #2138307, #2137603, and #2138296. The content is solely the responsibility of the authors and does not necessarily represent the official views of the NIGMS, NIH, or NSF. The authors declare no conflict of interest.

## Supplementary Information for

## A. APPENDIX A

### CONSTRUCTING SEMI-EMPIRICAL PHASE DIAGRAMS FROM TAIL FITTING

Here, we construct semi-empirical FLD phase diagrams for the three pairs of influenza antigens without using the Gaussian approximation. Instead, we fit the tails of the distribution of free energy differences in order to obtain more accurate boundary values from extreme value statistics. Finally, we show that downsampling the antibody dataset by 90% still yields tail fits which are predictive of the phase boundaries for the full antibody dataset.

#### 1. Experimental log dissociation constant distributions

First, in Figure S1 we reproduce the log *K*_*d*_ distributions (equivalent to the Δ*G* distributions) for the three influenza antigens, where each datapoint represents one of up to up to 65, 535 antibodies whose binding affinities to each antigen were measured by [31]. The distributions are directly reproduced from the dataset; not all antibodies have binding affinites reported (and are stored as NaN values), which is why each antigen has a different number of antibody datapoints. Notably, the Tite-seq technique [32] used by [31] was limited to log *K*_*d*_ values stronger (i.e. more negative) than ™6, so antibodies which bound more weakly (i.e. log *K*_*d*_ *≥ ™*6) were automatically assigned a value of ™6, resulting in large spikes in two of the antigens’ log dissociation constant distributions (Figure S1(a-b)). In Figure S2, the joint log *K*_*d*_ distributions of for pairs of antigens are reproduced, also from [31]. In each subplot, each data point, which represents a unique antiobody, was included only if a measured log *K*_*d*_ was reported for both of the two antigens.

#### 2. Log dissociation constant difference distributions

Next, we compute the differences between each antibody’s log *K*_*d*_ for pairs of antigens:

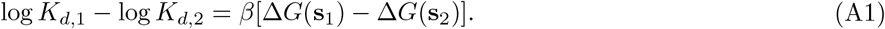

These difference distributions are plotted in Figure S3. As can be readily seen in the histograms, these distributions are non-Gaussian, so the approximation applied in the main text is not expected to hold necessarily. To obtain better estimates of phase diagram boundaries (relative to those obtained from the Gaussian approximation), we only need to capture the tail behaviors for each distribution.

#### 3. Extreme value theory for expected bound on distribution tails with finite samples

In extreme value statistics, the generalized Pareto distribution (GPD) is commonly used for this tail fitting task [40, 41], and it enables an analytically tractable estimate for the expectation of the tail’s maximum for a certain number of samples. The GPD’s probability density function is given by

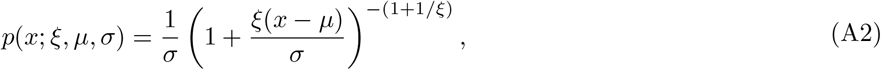

where *µ* is called the location parameter (*not* necessarily equal to the distribution’s mean), *σ* is the scale parameter, and *ξ* is the shape parameter. Following standard extreme value theory [40, 41], we first construct a cumulative distribution function for a random variable *X* = log *K*_*d*,1_ − log *K*_*d*,2_ which represents a datapoint drawn from the tail of the log dissociation constant difference distribution modeled by the GPD. The location parameter is set to *µ* = 0 because the distribution will be shifted through the introduction of a threshold. As is commonly done in extreme value theory, we shift the data distribution so that the tail begins at a threshold value *u*, which is moved to the origin. So, we define *Y* = *X* − *u*, and the CDF can be written

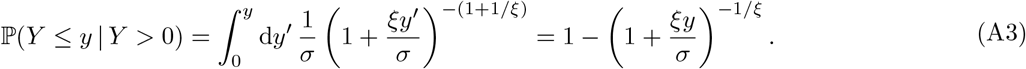

Undoing the affine shift, we can write

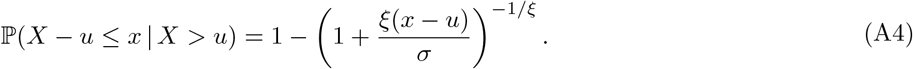

The probability of a datapoint being greater than a certain value *x within the tail* is given by the product of probabilities of drawing a datapoint from the tail the conditional probability above:

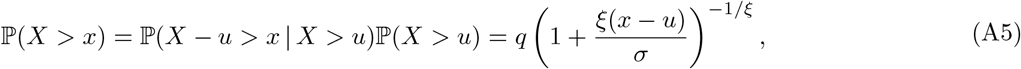

where we let *q ≡* ℙ (*X > u*) be a pre-defined threshold quantile to which fixes the start of the tail. Now, we define

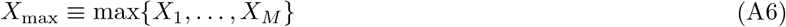

to be a random variable representing the maximum of *M* i.i.d. datapoints. The maximum will be less than some value *x* only if all drawn variables are also less than *x*:

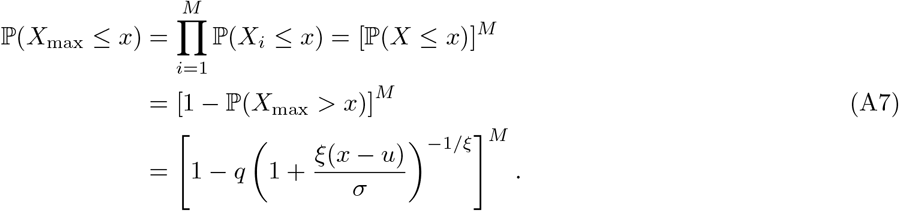

In the large *M* (many samples) limit, we take the typical Poisson limit commonly invoked for independent events:

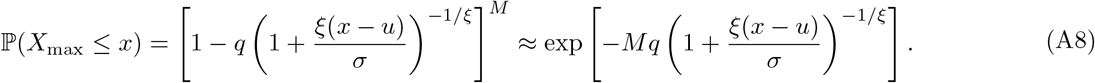

The CDF above defines a generalized extreme value (GEV) distribution from which we can quickly compute the expectation of the extremal value by integrating within the tail:

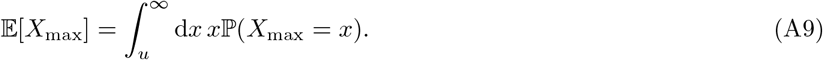

The PDF is computed by taking a derivative of the CDF with respect to *x*:

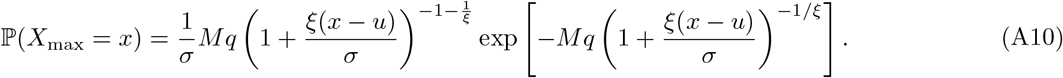

Performing the substitution

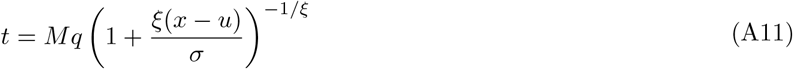

also gives us

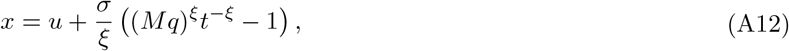

and

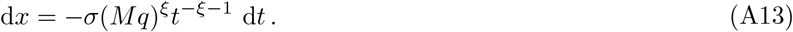

Now, the integral becomes

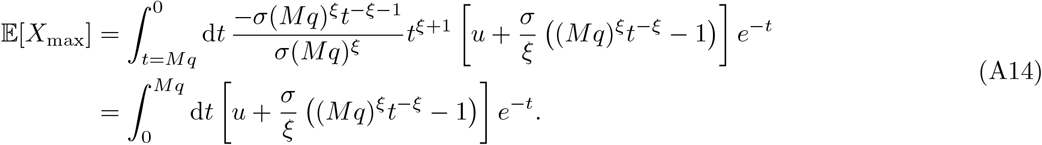

In the limit of large *M* (which we used in the Poisson limit earlier), we take *Mq* → ∞, so

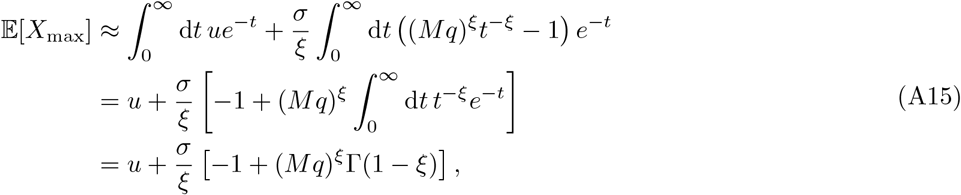

where we have used the Gamma function Γ and have assumed *ξ <* 1, which we will find later holds for all of the datasets studied here. By fitting the GPD PDF (eq. (A2)) with *µ* = 0 to the tails of our log *K*_*d*_ difference distributions, we can obtain estimates for *σ* and *ξ* and then substitute them into eq. (A15) in order to predict the expected bound on the data distribution for finite samples *M* . For an antibody library of size *M*, this is perhaps tautological, but below we will also show that we can downsample the antibody library to size 0.10 × *M* and still make predictions on the boundaries for the full library of size *M* .

#### 4. Fitting tails of log dissociation constant difference distributions

We now return to our empirical study of the log dissociation constant distributions. Although various methods exist for deciding where tail thresholds should be optimally placed [40, 41], we choose threshold quantiles arbitrarily to be the extremal 5% of the data distribution on either tail, *except* for the lower tail of the log *K*_*d*,SI06_ − log *K*_*d,G*189*E*_ distribution, for which we choose the bottom 20% in order to capture the large spike which is caused by the Tite-seq detection boundary.

After truncation based on the quantile thresholds described above, we reflect and/or affine shift the tails so that they are all in the non-negative real domain. Then, we fit the GPD to the tails using maximum likelihood estimation, as pre-implemented using SciPy’s <monospace>scipy.stats.genpareto.fit</monospace> [42]. From the estimated 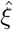 and 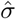 values, we calculated the expected bounds on the tails using eq. (A15). In Figure S4, we show the tails with the GPD fits, the estimated bounds from extreme value theory, and the actual experimentally observed bounds; the tails have been re-reflected and re-shifted to their original domains and orientations.

#### 5. Semi-empirical phase diagrams without the Gaussian approximation: tail fitting

Substituting the estimated bounds from eq. (A15) into main text eq. (9), we are able to construct semi-empirical phase diagrams (Figure S5) as an alternative to the phase diagrams computed from the Gaussian approximation in main text Figure 3. Compared to the Gaussian approximation, capturing the tail behaviors accurately yield excellent theoretical estimates. This is to be expected since we are essentially using the extremal data values from the empirical distribution to estimate the boundary, albeit somewhat indirectly through fitting *ξ* and *σ*.

A more powerful consequence of this tail fitting-based approach, however, is that *experimental scalability* is enabled, which we subsequently investigate by downsampling the antibody library.

#### 6. Scalability of phase boundary inference: downsampling the antibody library

We now downsample the antibody library to roughly 10% of its original size. We randomly choose 10% of the 65,535 indices; however, some antibodies have NaN reported as their log dissociation constant to particular antigens. So, the exact number of antibodies comprising the downsampled log dissociation constant difference distributions (Figure S6) is not exactly 10% of the full distributions from Figure S3. In general, the sample size of 10% captures the general shape of the histogram of the full distribution, but of course outliers are less likely to be sampled.

We then isolate tails using the same quantile cutoffs used for the full distribution, reflect and/or affine shift, and fit the GPD. Then, we calculate the expected phase diagram boundary using eq. (A15), but with *M* set to the *full* antibody dataset size (i.e. between 62, 000 and 65, 000, depending on how many NaN values were present). This means we are using the fitted parameters from the downsampled dataset but are *predicting* the phase boundaries for the full datset. Example tail fits from a single trial of the sampling procedure are shown in Figure S7. The downsampling procedure is conducted for 10, 000 trials, and the distributions for the expected extrema E[min(log *K*_*d*,1_ − log *K*_*d*,2_)] (for lower tails) or E[max(log *K*_*d*,1_ − log *K*_*d*,2_)] (for upper tails), constructed from those 10,000 trials, are shown in Figure S8, along with quartile boundaries, means, and the true extrema for comparison. The phase diagrams arising from the downsampling procedure are shown and discussed in main text Figure 4, with quartile boundaries plotted as well. The approach is repeated for 10,000 trials of downsampling the training set to size 1% × *M*, and the resulting phase diagrams are shown in Figure S9. Median boundaries are similar to those from the 10% datasets, but the phase boundary distributions are wider. The ability to use a smaller antibody sample to predict the phase boundaries for the full antibody library demonstrates the scalability of the experimental approach for elucidating FLD phase diagrams.

However, it should be noted that the GPD fits and predicted maxima are heavily dependent on outlier values. As we see in the main text, sometimes the phase boundary is underestimated if outliers are not captured in a particular sample. This is naturally always a limitation of extreme value prediction using smaller samples.

#### B. Supplementary Figures

**FIG. S1.**
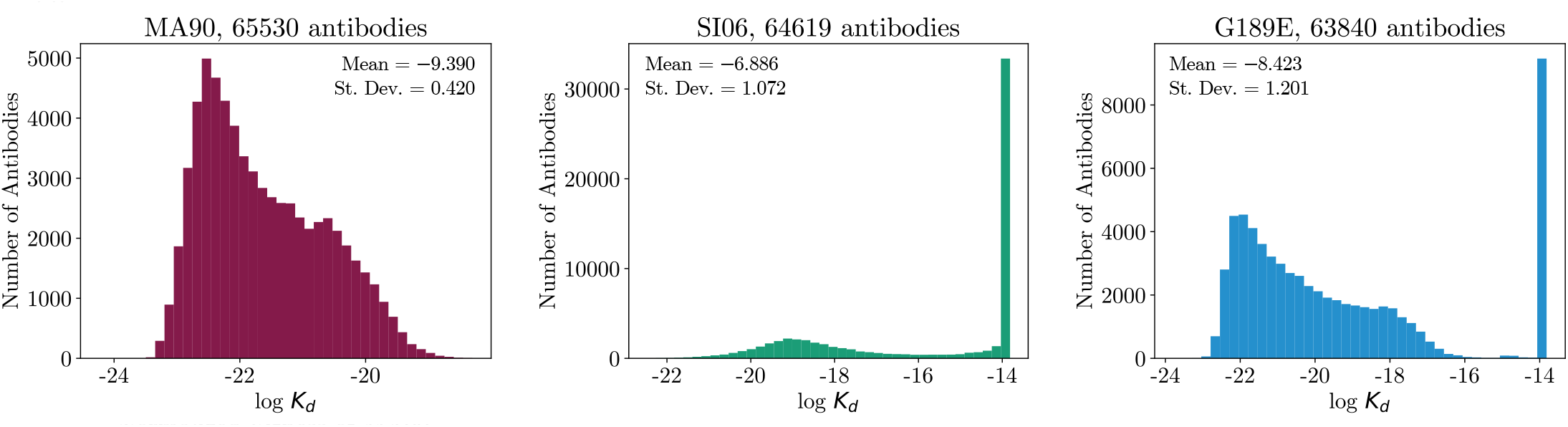
Distribution of log dissociation constants between over 62,000 antibodies and each of three influenza antigens, directly reproduced from the public dataset from [31]. Spikes in the distribution are due to weak binders whose log *K*_*d*_ ≥ −6 being binned to a value of −6.

**FIG. S2.**
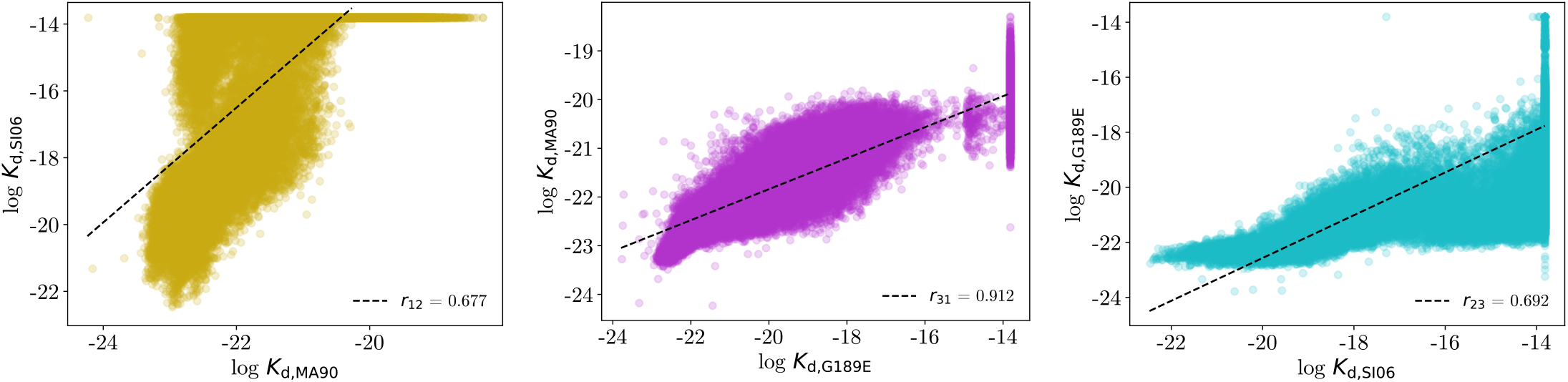
Joint distributions of log dissociation constants between over 62,000 antibodies and pairs of antigens, reproducing the data visualized in ref. [31], with Pearson correlations *r*12 shown. Spikes in the distribution are due to weak binders whose log *K*_*d*_ ≥ −6 being binned to a value of −6.

**FIG. S3.**
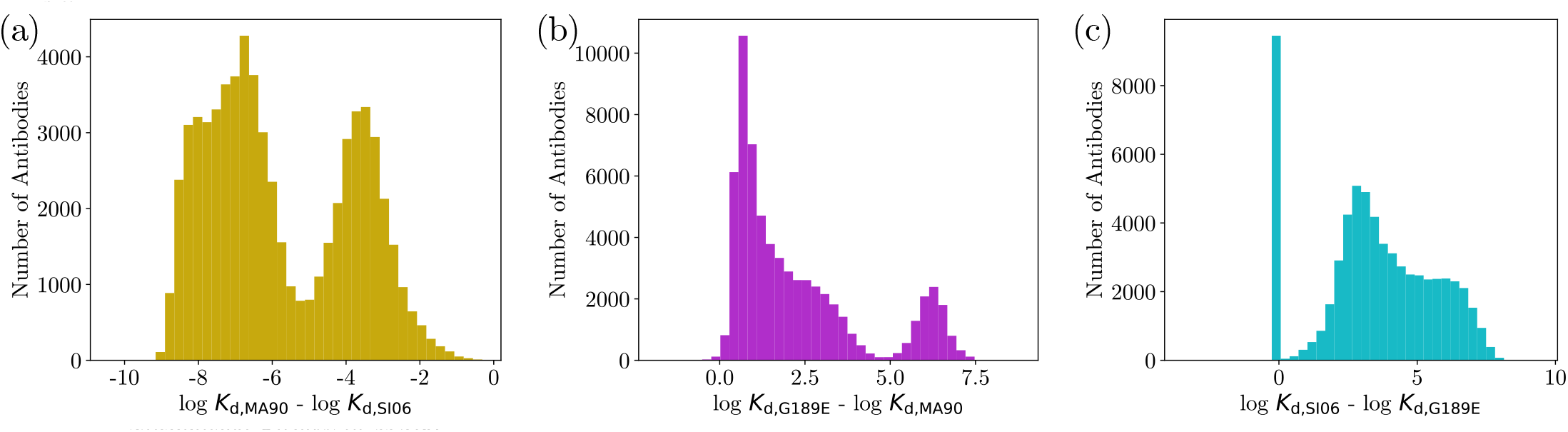
Distributions of the difference in log dissociation constant each antibody has to two different influenza antigens, as described by eq. (A1). Antibody count *M* differs for each distribution based on the number of NaN values in the dataset. (a) *M* = 64, 616, (b) *M* = 63, 835, and (c) *M* = 62, 929.

**FIG. S4.**
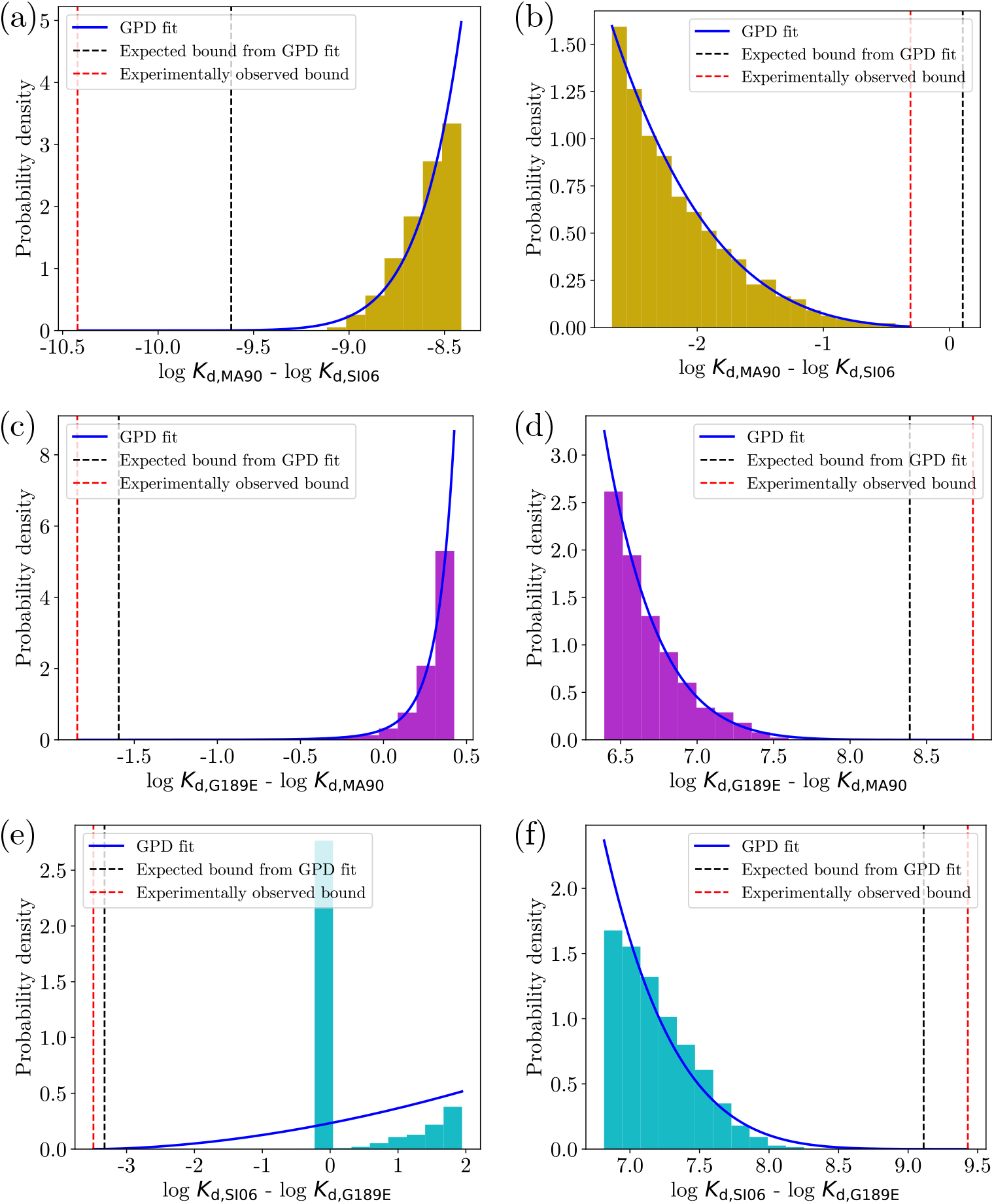
Upper and lower tails of the distributions in Figure S3 fit using the generalized Pareto distrbution in eq. (A2), with location parameter set to *µ* = 0. (a) Lower tail of Figure S1(a), (b) upper tail of Figure S1(a), (c) lower tail of Figure S1(b), (d) upper tail of Figure S1(b), (e) lower tail of Figure S1(c), (f) upper tail of Figure S1(c). 5% quantiles were used to threshold the tails for (a-d,f), and the lower 20% quantile was used to threshold the tail for (e), in order to capture the spike.

**FIG. S5.**
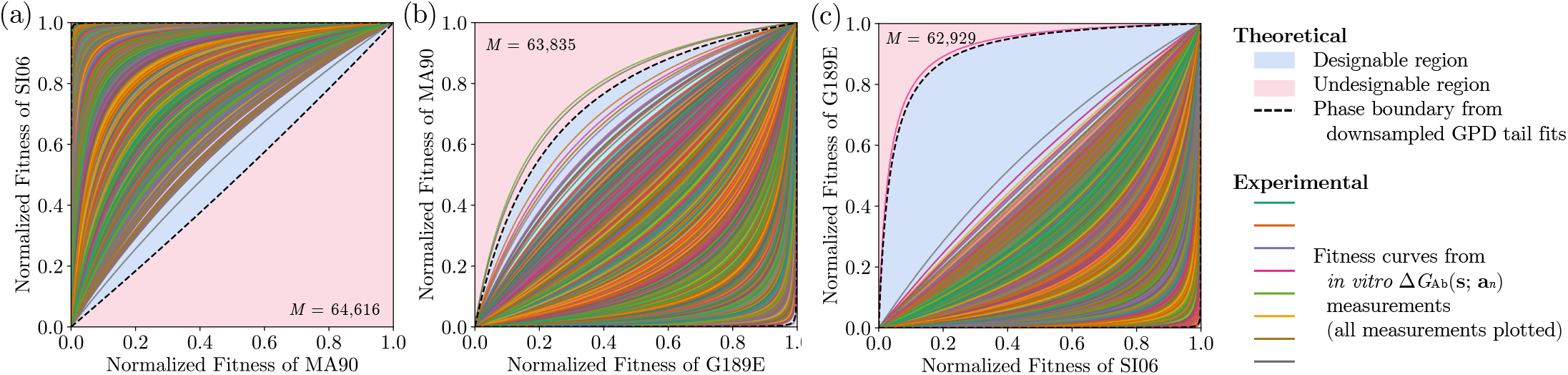
Semi-empirical phase diagrams constructed by substituting the estimated bounds on the log dissociation constant difference distributions (i.e. black dashed lines in fig. S4) into main text eq. (9). The theoretical phase boundaries show excellent agreement with the experimental extremal antibodies, as expected since the full data range is used to estimate the bounds.

**FIG. S6.**
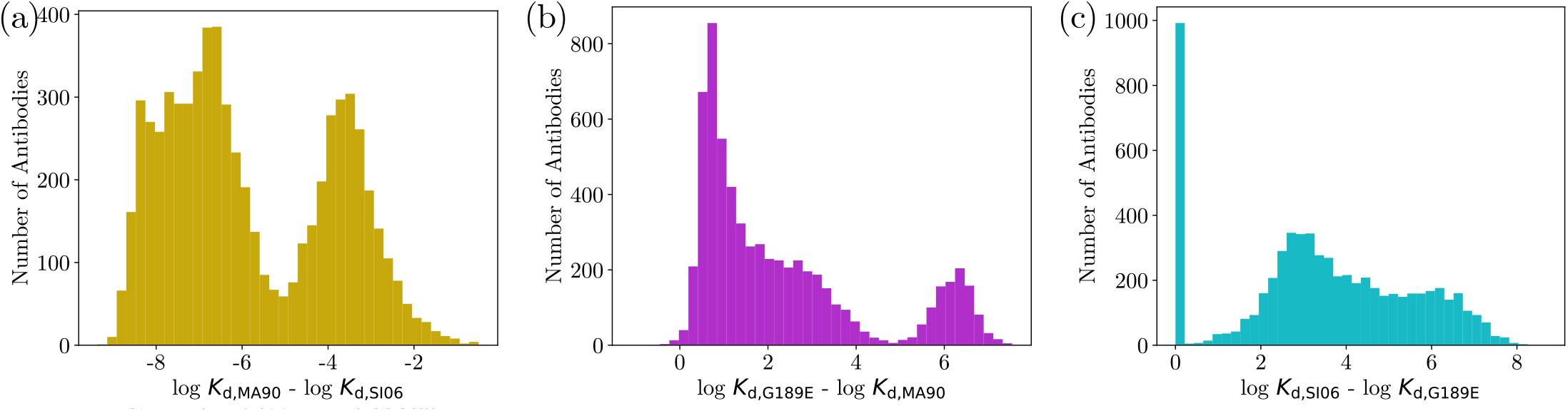
A representative example of log dissociation constant difference distributions for a single downsampling trial, using approximately 10% of the original antibody library. Antibody count *M* differs for each distribution (and may not be exactly 10% of the original datasets) based on the number of NaN values in the dataset. (a) *M* = 6, 469, (b) *M* = 6, 396, and (c) *M* = 6, 314.

**FIG. S7.**
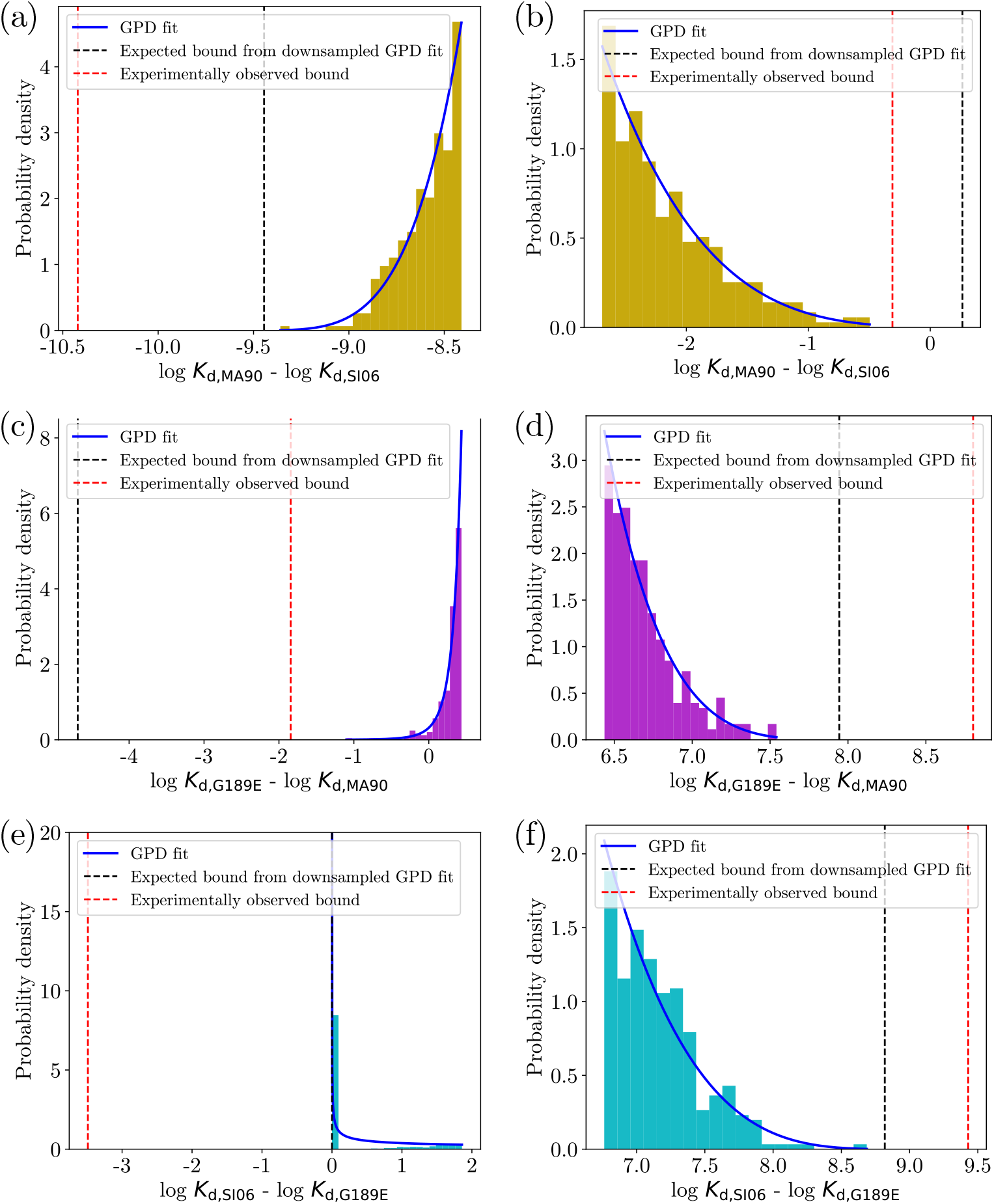
Upper and lower tails of the distributions in Figure S6 fit using the generalized Pareto distrbution in eq. (A2), with location parameter set to *µ* = 0. (a) Lower tail of Figure S6(a), (b) upper tail of Figure S6(a), (c) lower tail of Figure S6(b), (d) upper tail of Figure S6(b), (e) lower tail of Figure S6(c), (f) upper tail of Figure S6(c). 5% quantiles were used to threshold the tails for (a-d,f), and the lower 20% quantile was used to threshold the tail for (e), in order to capture the spike.

**FIG. S8.**
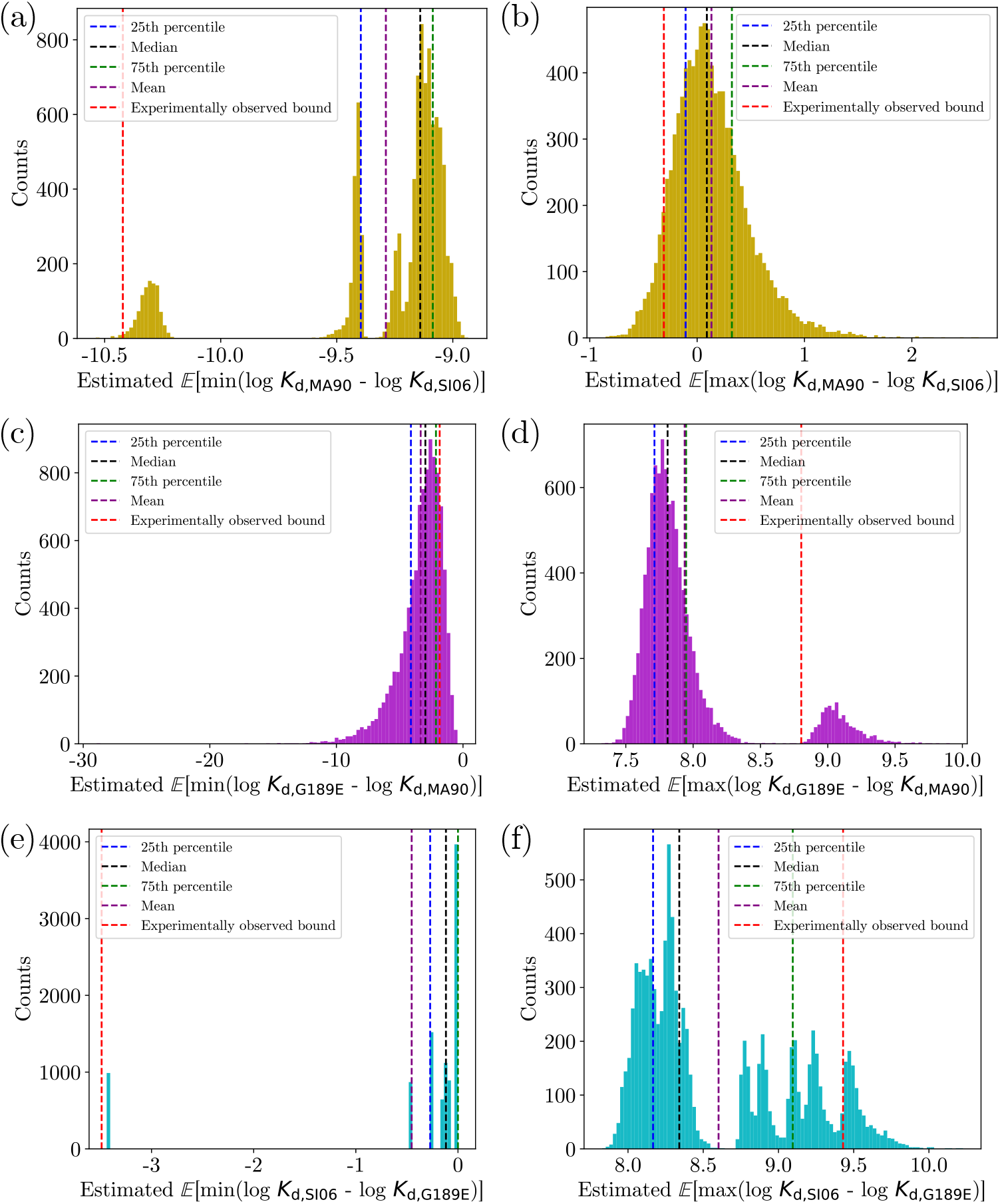
Distributions for the estimated extremal values for each tail, where each distribution has been built from 10,000 trials of the downsampling procedure. For lower tails, E[min(log *K*_*d*,1_ − log *K*_*d*,2_)] is calculated for antigens 1 and 2, and for upper tails E[max(log *K*_*d*,1_ − log *K*_*d*,2_)] is calculated for antigens 1 and 2. (a) Lower tail for MA90 vs. SI06, (b) upper tail for MA90 vs. SI06, (c) lower tail for G189E vs. MA90, (d) lower tail for G189E vs. MA90, (e) lower tail for SI06 vs. G189E (f) upper tail for SI06 vs. G189E. Vertical lines indicate 25th, 50th (i.e. median), and 75th percentiles as well as the empirical mean and the experimentally observed extrema.

**FIG. S9.**
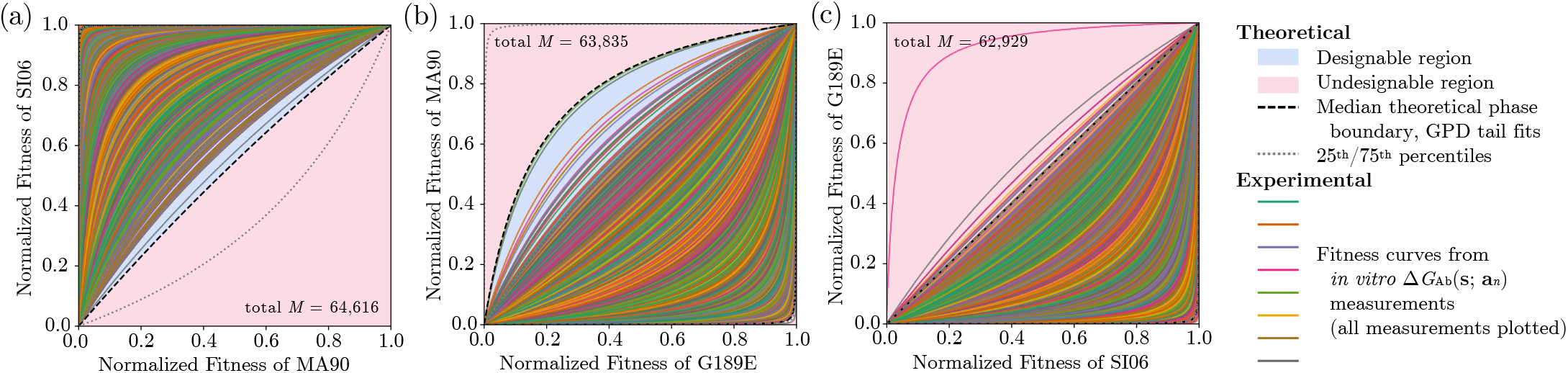
Experimental FLD phase diagrams from log dissociation constant measurements of over 62,000 antibodies with three influenza glycoprotein antigens, and semi-empiricial phase boundaries from fitting the tails of downsampled antibody datasets which are approximately 1% of the size of the original dataset. Empirical measurements are shown as solid, colorful lines. Estimated median phase boundaries from 10,000 independent downsampling trials are shown as black, dashed lines, and 25th/75th percentile phase boundaries are shown as gray, dotted lines.

